# Selective loss of the GABA_Aα1_ subunit from Purkinje cells is sufficient to induce a tremor phenotype

**DOI:** 10.1101/773655

**Authors:** Angela Nietz, Chris Krook-Magnuson, Haruna Gutierrez, Julia Klein, Clarke Sauve, Esther Krook-Magnuson

**Author notes:** **Corresponding Author:** [The authors have declared that no conflict of interest exists.].

## Abstract

**Background:** Increased circuit level insights into Essential tremor, the most prevalent movement disorder, are needed. Previously, an Essential Tremor-like phenotype was noted in animals with a global knockout of the GABA_Aα1_ subunit. However, global knockout of the GABA_Aα1_ subunit has limitations, including potential early mortality and limited circuit level insights into the tremor.

**Methods:** Given the hypothesized role of the cerebellum in tremor, including Essential Tremor, we used transgenic mice to selectively knock out the GABA_Aα1_ subunit from cerebellar Purkinje cells. As previous work suggested background strain may influence phenotype in this model, we used two different background strains (a Black6 and a Mixed background). We examined the resulting phenotype regarding impacts on inhibitory postsynaptic currents, survival rates, gross motor abilities, and expression of tremor.

**Results:** We found that GABA_A_-mediated synaptic currents are abolished in Purkinje cells from Purkinje cell specific knockout mice, while GABA_A_-mediated inhibition to cerebellar molecular layer interneurons remains intact. Selective loss of GABA_Aα1_ from Purkinje cells did not produce gross motor deficits, as measured by the accelerating rotarod, nor did it result in decreased survival rates. However, a tremor phenotype was apparent, regardless of sex or background strain. This tremor mimicked the tremor seen in animals with a global knockout of the GABA_Aα1_ subunit, and, like Essential Tremor in patients, was responsive to ethanol.

**Conclusions:** These findings indicate that reduced inhibition to Purkinje cells is sufficient to induce a tremor phenotype, highlighting the importance of the cerebellum, inhibition, and Purkinje cells, in tremor.

## INTRODUCTION

Essential Tremor is one of the most common, yet least understood movement disorders, with prevalence rates far surpassing Parkinson’s Disease (1–5). A major limitation in the study of Essential Tremor is a lack of sufficient animal models available to study the underlying brain circuitry, pharmacology behind anti-tremor drugs, and development of novel treatments (6). Mice with a global knockout of the GABA_Aα1_ subunit have been proposed as a model of Essential Tremor (7). The tremor observed in global GABA_Aα1_ knockout mice shows similar pharmacology to human Essential Tremor, including sensitivity to ethanol (7, 8). A role for dysfunction in GABAergic signaling in tremor is more broadly supported (9–13). However, as GABA_Aα1_ is expressed in a variety of cell types throughout the central nervous system, it is unclear which brain region(s) may be responsible for the tremor phenotype in global GABA_Aα1_ knockout mice.

The cerebellum has been implicated in Essential Tremor (9, 14–17) (but see also e.g. (18–23) regarding other important areas) and post-mortem studies have repeatedly found significant pathology in the cerebellum of Essential Tremor patients (1, 11, 13, 24–26). Similarly, tremor (rest, action, orthostatic, and intentional) phenotypes are noted in a range of disorders with significant cerebellar components, including spinocerebellar ataxias (SCAs) (27, 28). We hypothesized therefore that selective loss of GABA_Aα1_ from cerebellar Purkinje cells could recapitulate the tremor phenotype observed in global GABA_Aα1_ knockout animals. If a tremor phenotype is observable when GABA_Aα1_ is removed exclusively from Purkinje cells, a key role of the cerebellum in tremor would be firmly established, and other non-tremor phenotypes associated with the global knockout of GABA_Aα1_ may be avoided.

As subunit composition contributes to channel kinetics (29–31), removal of the GABA_Aα1_ subunit can alter the decay kinetics of GABA_A_-mediated inhibitory post-synaptic currents (IPSCs) (32, 33). Therefore, in the global GABA_Aα1_ knockout mice it is additionally unclear if the tremor phenotype arises from a deficit of inhibition or an alteration in decay kinetics (34). However, while most neurons express multiple isoforms of the α subunit, adult Purkinje cells appear to exclusively express the α1 isoform (35, 36). As all known functional GABA_A_ receptors contain an α subunit (37) (with the notable exception of ρ containing “GABA_C_” receptors), selective knockout of the GABA_Aα1_ subunit can result in complete loss of GABA_A_-mediated inhibition in Purkinje cells (36, 38, 39). In this case, if a tremor phenotype is observed in animals with a selective knockout of the GABA_Aα1_ subunit from Purkinje cells, it will be attributable specifically to a deficit of inhibition, rather than a change in the kinetics of remaining inhibition. This would provide further insight into the pathophysiology underlying tremor.

We produced mice with a selective knockout of GABA_Aα1_ from Purkinje cells. Previous studies have suggested that the phenotype displayed in the global knockout model may depend on the background strain of the mice (40). Therefore, we examined the resulting phenotype in Purkinje cell specific GABA_Aα1_ knockout mice on both a Mixed and Black6 genetic background. We find that a tremor phenotype is present in animals with a knockout of GABA_Aα1_ selectively from Purkinje cells, regardless of the background of the animal, firmly establishing the relevance of the cerebellum (and inhibition to Purkinje cells in particular) to tremor.

## METHODS

All experimental protocols were approved by the University of Minnesota’s Institutional Animal Care and Use Committee.

### Animals

To selectively knockout GABA_Aα1_ expression from Purkinje cells, we crossed floxed-GABA_Aα1_ mice (B6.129(FVB)-Gabra1tm1Geh/J; Jackson stock number: 004318) (7, 41) with mice expressing Cre-recombinase in Purkinje cells (Pcp2-Cre mice; B6.Cg-Tg(Pcp2-cre)3555Jdhu/J; Jackson stock number: 010536) (42, 43). While Cre expression is also found in retinal bipolar neurons in Pcp2-Cre animals, this Pcp2-Cre line does not show widespread leaky expression (43, 44). Purkinje cell expression of Cre starts within the first postnatal week (45), resulting in gradual loss of GABA_Aα1_ immunoreactivity from Purkinje cells during the second and third postnatal week (46), after the establishment of climbing fiber one-to-one innervation of Purkinje cells (47).

As the phenotype of the global GABA_Aα1_ knock-out mouse has been suggested to be dependent on the background of the animals (7, 40), we produced mice with a selective knockout of GABA_Aα1_ on both a Black6 background (“PC-KO-Bl6” mice) and a mixed background (“PC-KO-Mixed”). To obtain breeders on a Mixed background, we crossed floxed-GABA_Aα1_ breeders with both FVB/NJ (Jackson stock number 001800) and 129S1/SvlmJ (Jackson stock number 002448) mice.

Animals homozygous for WT GABA_Aα1_ and/or Cre negative were considered “PC-WT”. Animals which were Cre positive and hemizygous for floxed-GABA_Aα1_ were considered “PC-HEMI”, and animals which were Cre positive and homozygous for floxed-GABA_Aα1_ were considered “PC-KO”.

Animals with a global knockout of GABA_Aα1_ (“Global-KO”) were also generated, allowing a direct comparison of tremor phenotypes in this study. In order to mimic previous work examining tremor in global knockout animals, we generated Global-KO animals on a Mixed background. First, floxed-GABA_Aα1_ mice were crossed with FVB/NJ and 129S1/SvlmJ mice as described above. These offspring were then crossed with mice broadly expressing Cre recombinase (B6.C-Tg(CMV-cre)1Cgn/J; Jackson stock number: 006054; note that this includes germline expression, such that continued expression of Cre is not necessary). Hemizygous mice were then crossed, and Cre removed in subsequent litters by selective breeding.

Both male and female animals were used in experiments. Where found, sex differences are noted in the Results section.

### Slice electrophysiology

Mice (23-392 days old) were anesthetized with isoflurane inhalation followed by intraperitoneal (i.p.) injection of 2, 2, 2-tribromoethanol or by 2, 2, 2-tribromoethanol injection alone and intra-cardially perfused with ice cold cutting solution before decapitation. The brain was removed from the skull, and the cerebellum was dissected in ice cold sucrose cutting solution containing (in mM): 85 NaCl, 75 Sucrose, 2.5 KCl, 25 Glucose, 1.25 NaH2PO_4_, 4 MgCl_2_, 0.5 CaCl_2_, and 24 NaHCO_3_. Sagittal slices of the cerebellum (300 μm) were cut and incubated for 30 min at 35ºC in the same sucrose cutting solution and subsequently incubated at room temperature until recordings. Recordings were performed in artificial cerebrospinal fluid (ACSF) containing (in mM): 2.5 KCl, 10 Glucose, 126 NaCl, 1.25 NaH2PO_4_, 2 MgCl_2_, 2 CaCl_2_, and 26 NaHCO_3_, at ~32ºC. Pipettes (2-5MΩ) were filled with an intracellular solution consisting of (in mM): 110 CsCl, 35 CsF, 10 Hepes, 10 EGTA.

Inhibitory post-synaptic currents (IPSCs) were recorded with a holding potential of −60mV and in the presence of the AMPAR and NMDAR antagonists NBQX (5µM; Tocris, #0373) and APV (10µM; Tocris, #0106). In a subset of cells, zolpidem (1µM; Sigma, #Z103) -- a GABA_Aα1_ subunit selective positive modulator -- was bath applied to assay the subunit composition of GABA_A_ receptors mediating IPSCs. To confirm IPSCs were due to GABA_A_ receptor activation, the GABA_A_ receptor antagonist gabazine (5µM; Sigma, #S106) was bath applied at the end of the experiment. IPCSs were analyzed using Clampfit 10.7, including its event detection feature. A weighted τ was calculated per cell on an averaged IPSC according to the equation 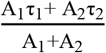, where A is the amplitude of each component and τ is the associated time constant.

### Kaplan-Meier survival curves

In order to examine if PC-KO animals displayed any early mortality, detailed record keeping was maintained, noting the date and cause of death for each animal. Animals euthanized due to health concerns were considered dead at the time of euthanasia. Animals euthanized solely for experimental needs (e.g., electrophysiology), or because the animal was no longer needed for experiments, were counted as censored observations. Animals still alive at the time of calculation of survival curves had the date of calculation as their final observation point (censored time point).

### Accelerating rotarod

Accelerating rotarod testing and analysis was performed utilizing resources of the Mouse Behavior Core, University of Minnesota. The experimenter was blinded to the genotypes. The rotarod apparatus had a diameter of 3 cm (Ugo Basile, #47600) and was set to a minimum speed of 5 revolutions per minute (RPM) and a maximum speed of 50 RPM, with an acceleration rate of 9 RPM per minute. A trial was deemed complete when the animal fell onto the paddle below or made two consecutive rotations without recovery of stepping. Animals were run for 4 trials a day with 15-20 minutes between trials. During the ‘training period’, animals were run on the task for 4 consecutive days (4 trials a day). During the ‘testing period’, mice were tested once a week for 5 weeks (4 trials a day) to assess retention on the task.

### Tail suspension

Tremor was measured using a custom-made tail suspension apparatus. The measurement sensor was a miniature load cell (force transducer: Honeywell, Golden Valley, MN, model 31) attached to an in-line amplifier (Sensotec Inc., Columbus, OH, part number: 008-0296-00). Signals were digitized (DAQ, National Instruments USB-6221) and recorded using custom Matlab-based software. A rotating attachment allowed the animals to be secured by the tail to the transducer while minimizing the risk of torque-induced damage to the transducer. Except when testing the pharmacological sensitivity of the tremor, trials lasted ten minutes, with data averaged across trials per animal. When testing the pharmacological sensitivity of the tremor, an initial, baseline, trial of 5 minute tail suspension was followed by an i.p. injection of either 1.25g/kg ethanol (at 10mL/kg) or saline (also 10mL/kg). Tail suspension trials (5min duration each) were then completed 20min, 40min, and 60 minutes after i.p. injection in order to assess the impact of ethanol administration on the recorded tremor. Habituation, saline, and ethanol experiments were run on separate days, with at least one day between experiments.

For each file, body weight of the animal was calculated based on the average recorded signal; these values closely matched weights manually collected and recorded. Recorded signals were then converted into percent change in force (relative to the mean force, i.e., weight). A “Tremor Metric” (TM) was calculated by taking the ratio of the power in the tremor range (20-30Hz) to the power in the adjacent ranges (15-20Hz and 30-35Hz). Power in each band was calculated using the matlab function *bandpower*. Moving time spectrograms, power spectral density (PSD) plots, and detailed analysis of bouts of tremor were done using the Chronux (http://chronux.org/) library (48).

We used the following operational definition of a tremor bout: the maximum power in the tremor band (20-30Hz) exceeds a threshold of 10^-4.5 *and* the portion of power (from 0-50Hz) present in the tremor band (i.e., 20-30Hz) exceeds a threshold of 0.4 (40%); these thresholds were determined empirically using datasets with clear tremor envelopes. Initial detections were combined if the gap separating the detections was less than 0.1s. The length of bouts was calculated with the start of the bout defined as the moment the enveloped signal (using matlab command *envelope*) first grew larger than a threshold of 9% change in force (relative to mean force), and the end as the moment it dropped below that threshold. Flagged ‘bouts’ with a duration less than 0.15s were removed, and not further considered in analyses.

### Statistical Analysis Software Used

Data sets were analyzed using Excel, Clampfit 10.7, OriginPro, and Matlab, including the Chronux (http://chronux.org/) library (48).

## RESULTS

### Selective knockout of the GABA_Aα1_ subunit from Purkinje Cells causes a loss of synaptic inhibition in Purkinje Cells

To confirm that our approach resulted in selective knockout of the GABA_Aα1_ subunit from Purkinje cells, we performed whole-cell patch-clamp recordings of IPSCs from Purkinje cells in cerebellar slices from PC-KO, PC-WT, and PC-HEMI mice (Figure 1).

**Figure 1.**
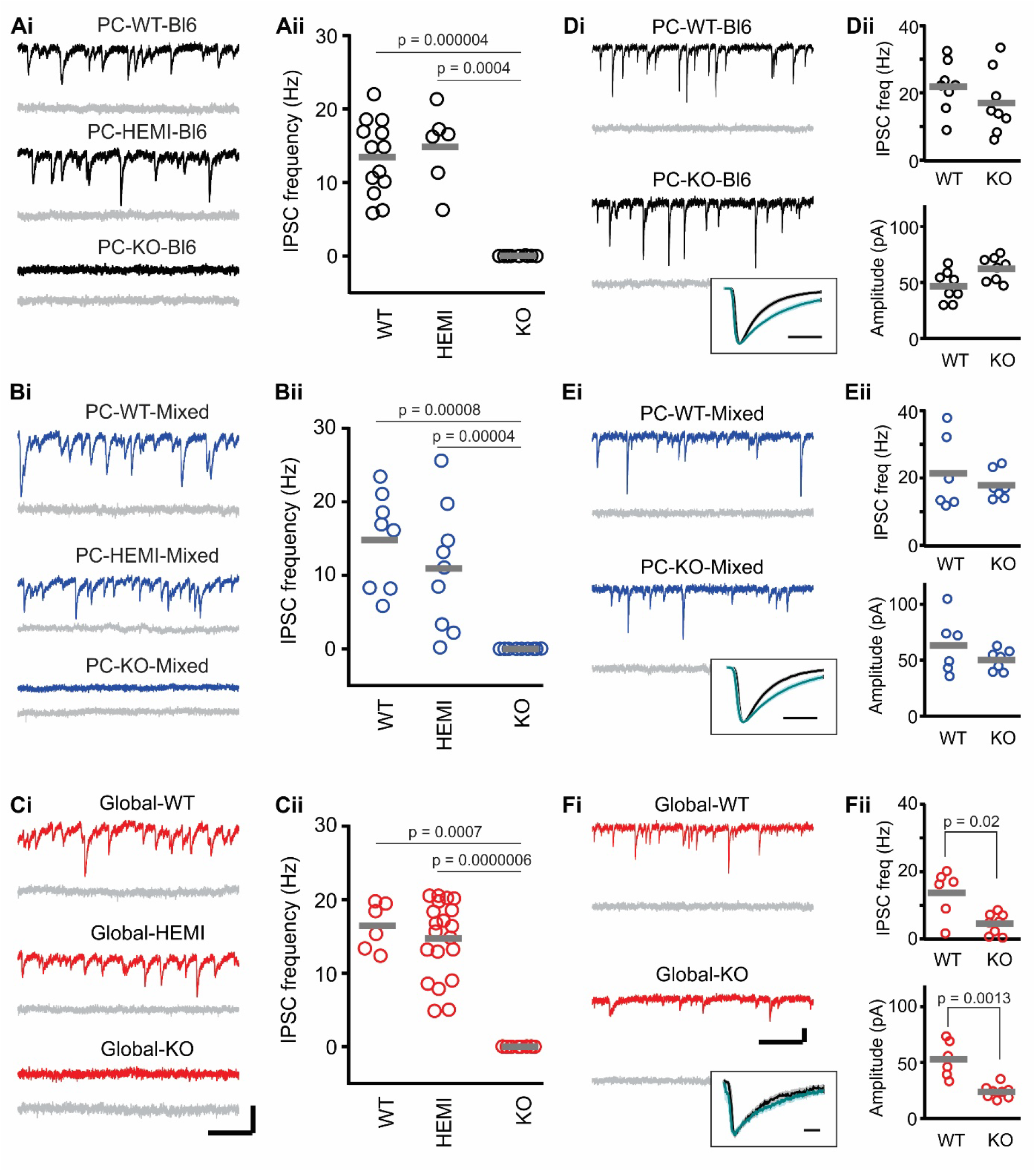
In PC-KO mice, GABA_A_-mediated IPSCs are selectively lost in Purkinje cells. Spontaneous IPSCs recorded from Purkinje cells (subsequently blocked by gabazine, grey traces) in PC-Bl6 (**A**), PC-Mixed (**B**), and Global (**C**) WT, HEMI, and KO animals. **D-F)** IPCSs recorded from molecular layer interneurons. **Insets:** Average molecular layer interneuron IPSCs (normalized amplitude) prior to (black) or in the presence of zolpidem (teal traces) for KO animals. Light shading indicates SEM. Note that IPSCs are lost in Purkinje cells in KO animals, but are unaffected in PC-KO molecular layer interneurons. (ii) Grey bars indicate mean. Scale bars: 100 pA, 200ms (A-C), 50 pA,200 (D-F), 5 ms (insets). See also Table 1.

PC-HEMI mice, which still express one functional copy of the GABA_Aα1_ gene, displayed a frequency of IPSCs which was not statistically different from WT animals (Table 1; Fig.1A-B). Similarly, amplitude and decay kinetics were not statistically different between PC-HEMI and PC-WT animals (Table 1). This suggests that expression levels of GABA_Aα1_ are adjusted in hemizygous mice to maintain an appropriate level of synaptic inhibition.

**Table 1.**
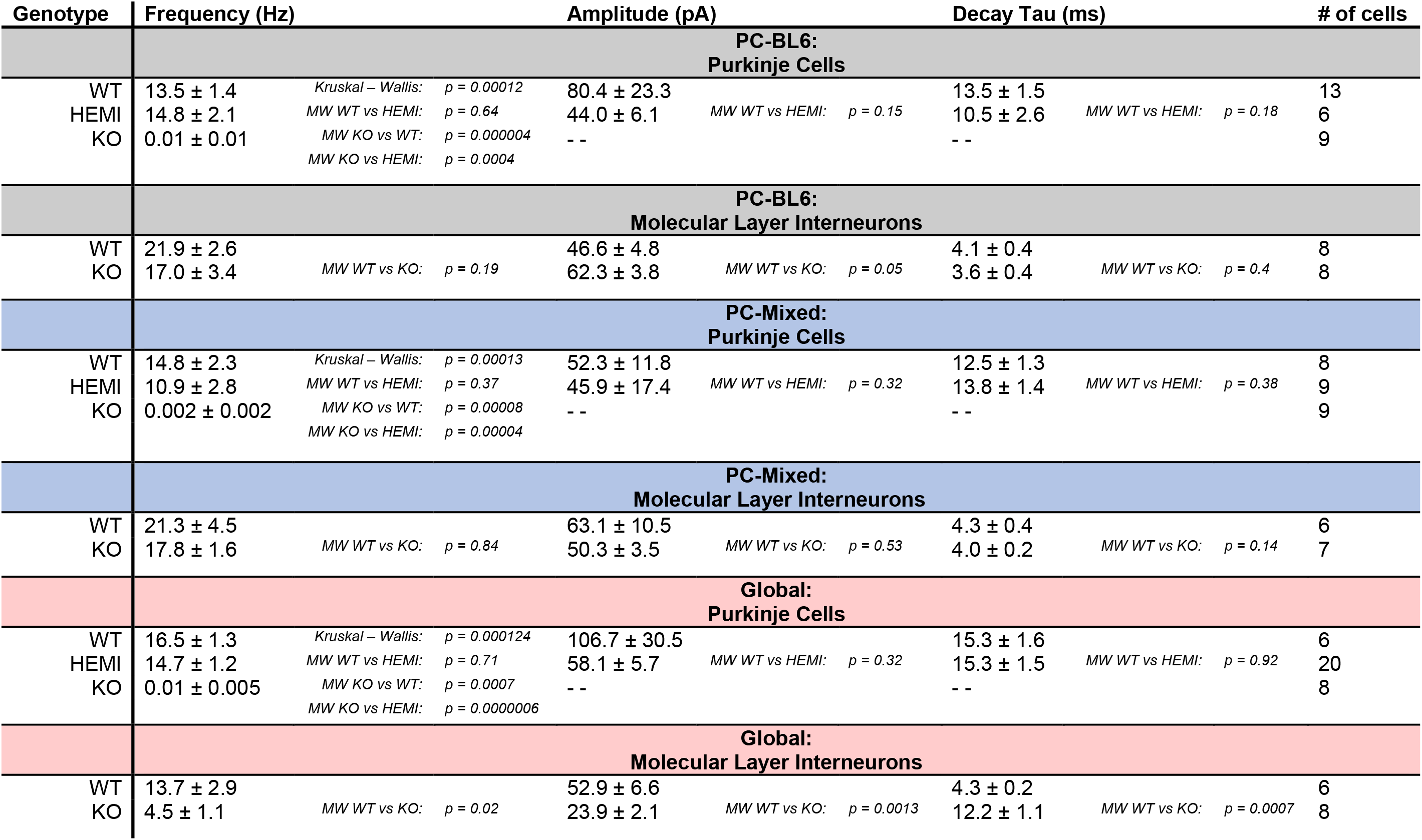
Properties of Purkinje cell and Molecular Layer Interneuron IPSCs. Average (±SEM) values and statistical comparisons of IPSCs by cell type and mouse line. Note that no values are given for amplitude or decay for Purkinje cells from KO animals, and one cell was excluded from PC-Mixed-HEMI decay analysis, due to too few IPSCs.

In contrast, Purkinje cells from PC-KO animals displayed a loss of IPSCs (Table 1; Fig.1A-B). These findings support previous work indicating that the α1 subunit is the only α subunit expressed in adult Purkinje cells, and loss of the α1 subunit results in a loss of all GABA_A_-mediated synaptic inhibition in Purkinje cells, without a compensatory increase in the expression of a different GABA_Aα_ subunit (36, 38, 39). For comparison, we also examined IPSCs in animals with a global, rather than Purkinje cell specific, knockout of GABA_Aα1_ (“Global-KO”). As expected, similar to PC-KO animals, global knockout of the GABA_Aα1_ subunit resulted in a loss of IPSCs in Purkinje cells (Table 1; Fig 1C). These data show our approach is able to knockout the α1 subunit from Purkinje cells, which results in a loss of synaptic GABAergic IPSCs in Purkinje cells.

### Selective knockout of GABA_Aα1_ from Purkinje Cells leaves inhibition to other cells intact

To check the specificity of our Purkinje cell specific GABA_Aα1_ knockout, we recorded IPSCs from cerebellar molecular layer interneurons which should not be impacted in Purkinje cell selective knockout animals. As anticipated, IPSCs recorded from molecular layer interneurons displayed similar amplitude, decay kinetics, and frequency in PC-WT and PC-KO animals (Table 1, Fig.1D-E). In a subset of molecular layer interneurons, we applied zolpidem, a positive allosteric modulator of GABA_A_ receptors containing the α1 subunit (49), to further confirm the presence of the GABA_Aα1_ subunit. As expected, we found a consistent increase in the decay tau after the application of 1µM zolpidem (weighted τ pre-zolpidem: 4.2 ± 0.2ms, vs post-zolpidem: 8.6 ± 1.0 ms, n = 11 molecular layer interneurons, p = 0.001, Wilcoxon, Fig 1D-E **insets**), further confirming the expression of the GABA_Aα1_ subunit in molecular layer interneurons in PC-KO animals.

We also examined IPSCs in molecular layer interneurons in Global-KO animals, as done previously by Vicini and colleagues (41), and confirmed that the frequency and amplitude of IPSCs was significantly reduced (Table 1, Fig. 1F). Moreover, due to lack of α1 expression, the decay kinetics of IPSCs in molecular layer interneurons were altered in Global-KO animals (Table 1) and were insensitive to zolpidem (n=6 cells, p=0.16 Wilcoxon, Fig. 1F **inset**). These results are in contrast to IPSCs in molecular layer interneurons in PC-KO animals and further affirms the sensitivity of our methods.

Therefore, in Global-KO animals, observed phenotypes may be due to effects on Purkinje cells *and/or* other cell types, and due deficits of inhibition (in the case of Purkinje cells) *and/or* changes in kinetics of inhibition (e.g. in molecular layer interneurons). In contrast, phenotypes observed in PC-KO animals can be attributed, specifically, to loss of inhibition to Purkinje cells.

### Selective knockout of GABA_Aα1_ from Purkinje Cells does not reduce survival rates

Having confirmed the specificity of our selective knockout approach, we next sought to examine resulting phenotypes. Mice with a global knockout of the GABA_Aα1_ subunit can display early death (40, 50). We also observed this early mortality in our hands (p<0.0001, Log Rank, comparing survival rates between genotypes in the global knockout line). Early mortality in Global-KO animals was concentrated around the time of weaning (Figure 2A), with higher rates of mortality in males (Fig. 2B). There was also, as previously reported (40), a decrease in body weight among Global-KO animals (WT vs HEMI: p=0.88, KO vs WT: p<0.0001, KO vs HEMI: p<0.0001, F-test).

**Figure 2.**
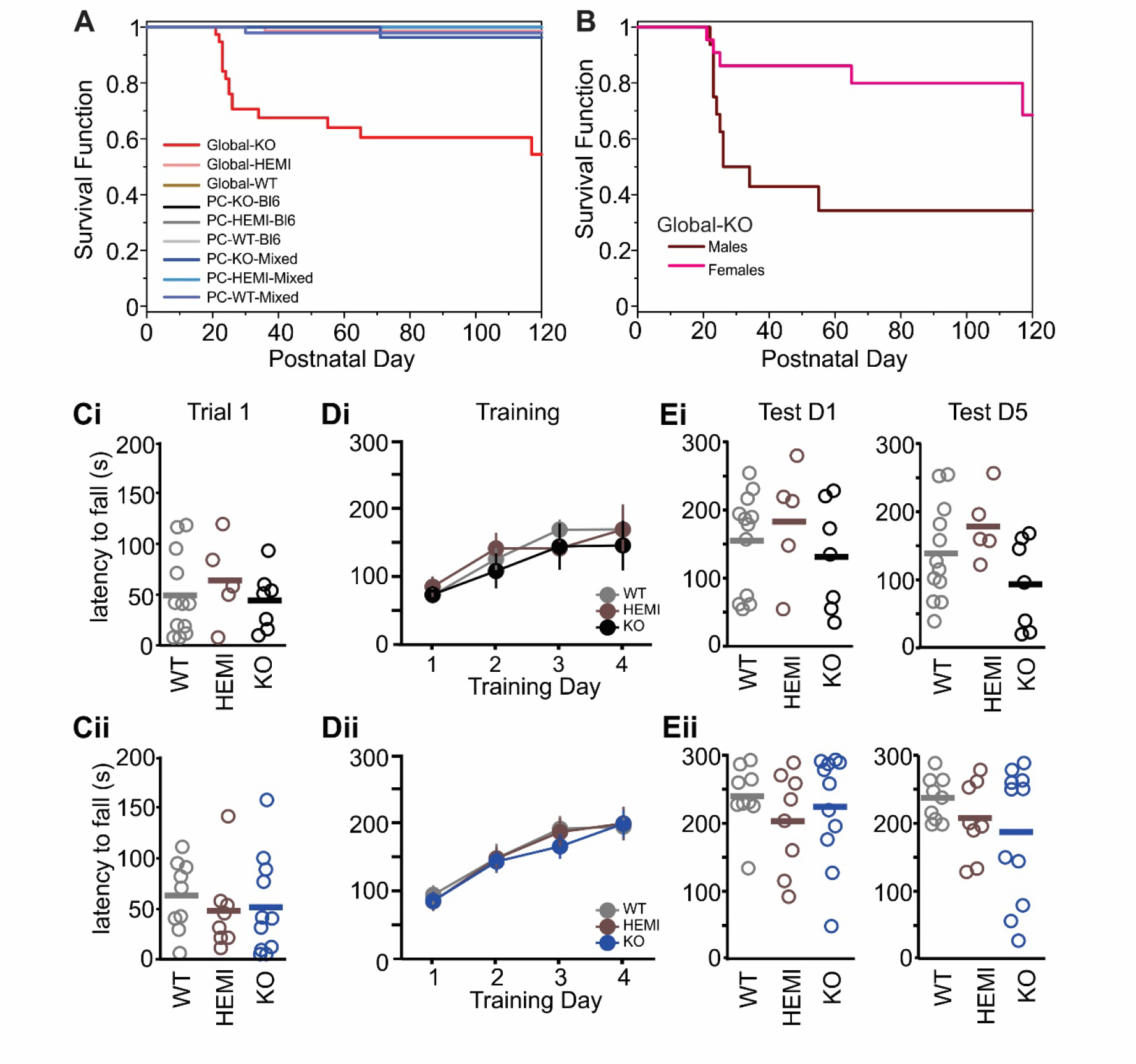
PC-KOs do not show reduced survival or impaired gross motor performance. (**A**) Kaplan-Meier survival curves show decreased survival rates in Global-KO animals (red trace), especially around the time of weaning (p<0.0001 across genotypes and mouse lines, Log Rank). Note that PC-KO animals show no increased mortality, and therefore lines overlap. (**B**) Decreased survival in Global-KO animals is especially pronounced in male animals (maroon line: male; pink line: female; male versus female p=0.01, Log Rank). (**C – E**) PC-KOs show no impairment on the accelerating rotarod task: mean latency to fall from the rotarod for the first trial of the first training day (**C**), across the four day training period (**D**), and on the first testing day (one week post-training, **E left**) and last testing day (five weeks post-training, **E right**) by genotype in PC-WT, PC-HEMI, and PC-KO animals on a Bl6 background (**i**) or Mixed background (**ii**). Bars: mean.

In contrast, PC-KO mice were generated at the expected Mendelian rates, and there were no differences in survival rates between PC-KO, PC-HEMI, and PC-WT animals, for either background (Fig. 2A). These data indicate that the decreased survival in Global-KO animals is not due to the loss of GABA_Aα1_ from Purkinje cells.

### Selective knockout of GABA_Aα1_ from Purkinje Cells does not produce gross motor deficits

As an initial assessment of gross motor abilities in the PC-KO animals, we utilized the accelerating rotarod test (Figure 2C-E). This allowed us to examine both baseline performance (by examining the first trial of the first testing day) as well as motor learning (by examining performance across training days) and retention (by examining performance across testing days). We found no significant differences between PC-KO, PC-HEMI, and PC-WT mice at baseline performance, regardless of background strain (Training day 1, trial 1, latency to fall: PC-WT-Bl6: 47 ± 11.8s, PC-HEMI-Bl6: 62 ± 18.4s, PC-KO-Bl6: 42 ± 11.0s, p=0.71 Kruskal-Wallis [KW]; PC-WT-Mixed: 62 ± 11.6s, PC-HEMI-Mixed: 47 ± 14.4s, PC-KO-Mixed: 51 ± 14.6s, p=0.55 KW; Fig. 2C), indicating similar initial motor performance. We also found no difference between PC-KO, PC-HEMI, and PC-WT mice, regardless of background strain, during training (training day * genotype: Black6 background p=0.83, Mixed background p=0.72; two way mixed effects ANOVAs; Fig.2D), indicating similar levels of motor learning across training trials. Finally, we also found no difference in performance on testing days (first test day: Black6 background p=0.62 KW, Mixed background p=0.59 KW; last test day: Black6 background p=0.14 KW, Mixed background p=0.51 KW; Fig.2E), suggesting similar retention of motor learning.

### Selective knockout of GABA_Aα1_ from Purkinje Cells results in a tremor phenotype

In global GABA_Aα1_ subunit knockout mice a tremor phenotype has been observed when animals are suspended by their tail (7, 51, 52). We therefore tail-suspended Global-KO animals while using a force-transducer to measure tremor, to confirm our ability to reliably detect tremor and provide a point of comparison for observations in PC-KO animals. Matching previous literature, we observed a tremor phenotype in our Global-KO animals when tail-suspended. This tremor was evident between 20Hz and 30Hz (Figure 3), as previously described for Global-KO animals (51, 52). Global-KO, but not Global-HEMI, animals displayed a tremor phenotype relative to Global-WT animals (relative power in the tremor band: “TM” Global-KO: 3.5 ± 0.3, n = 27 animals; TM Global-HEMI: 0.97 ± 0.03, n = 53 animals; TM Global-WT: 0.91 ± 0.04, n = 20 animals; p<0.0001 KW; WT vs HEMI p=0.25, WT vs KO p < 0.0001, HEMI vs KO p < 0.0001, post-hoc Mann-Whitneys [MWs]). We detected no difference in tremor severity between male and female Global-KO animals (TM male: 3.7 ± 0.79, female: 3.5 ± 0.26, p = 0.78 MW, Fig.3J).

**Figure 3.**
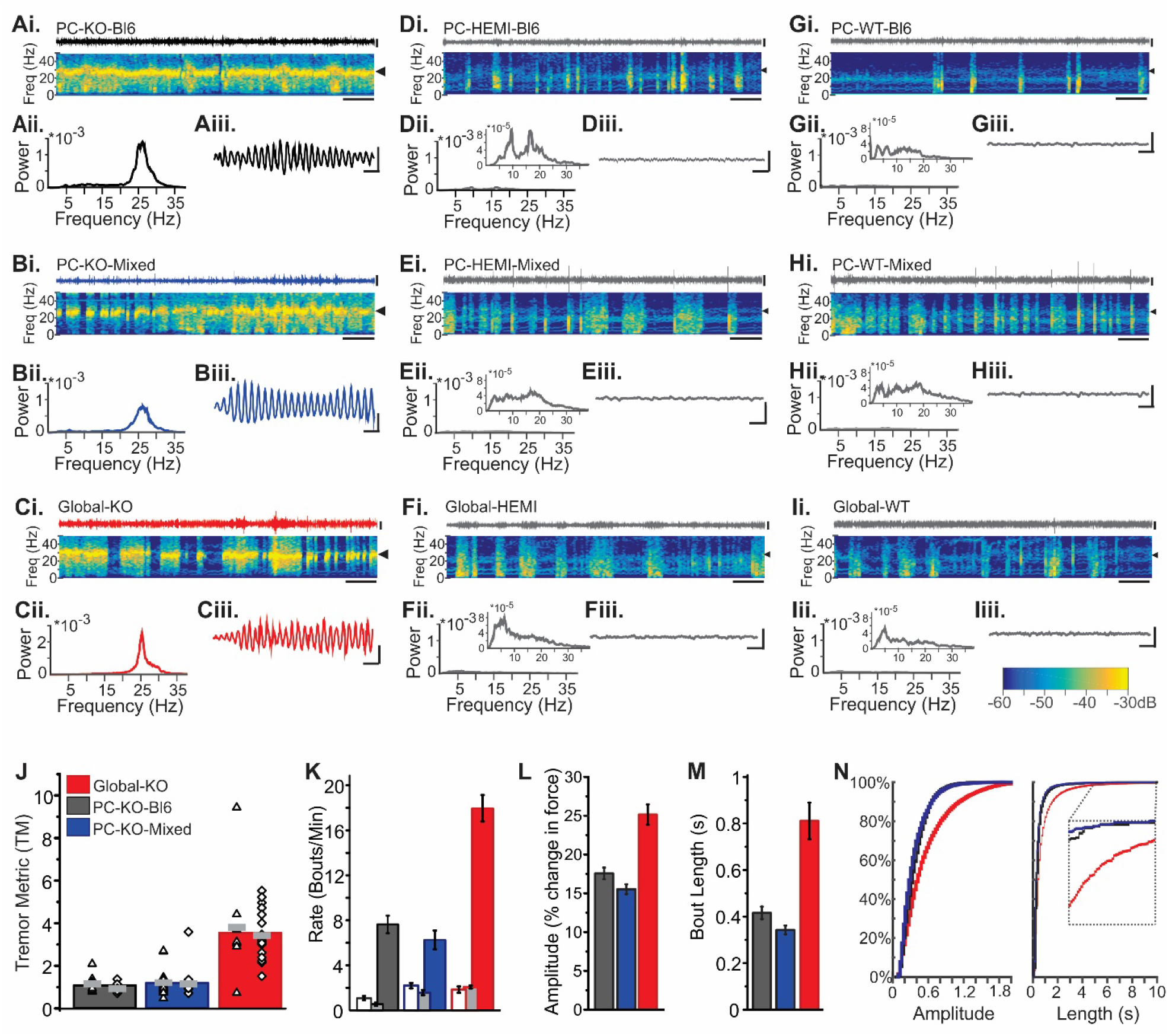
Selective knockout of GABA_Aα1_ from Purkinje cells is sufficient to produce a tremor phenotype. (**A-C**): (**i**) Top trace, example force recording for a PC-KO-Bl6 (**A**), PC-KO-Mixed (**B**) and a Global-KO (**C**) animal. Bottom: Corresponding spectrogram. Arrow highlights the tremor band. (**ii**) Corresponding power spectral density plots (PSDs) for each illustrated trace. (**iii**) One second traces illustrate recorded tremor oscillations. Similar data for HEMI (**D-F**) and WT (**G-I**) animals of each mouse line. Insets in **ii** for **D-I** have altered y-axes to allow visualization of the (lower amplitude) power spectrum in animals without a tremor. Color scale bar bottom right applies to all spectrograms. Scale bars **i**: 20s, 50% change in force; **iii**: 100ms, 50% change in force. (**J**) Male (triangles) and female (diamonds) animals show similar levels of tremor (grey lines represent mean for each sex) for a given mouse line. However, Global-KO animals (red bar) show greater levels of tremor compared to PC-KO animals of either background. (**K**) An automated analysis of bouts of tremor indicates a higher rate of tremor bouts in Global-KO animals compared to PC-KO animals. Open and light gray bars indicate the rate of detected tremor bouts in WT and HEMI animals, respectively, which provide an estimate of the rate of false bout detections (i.e. false positives). (**L-M**) Individual bouts were longer and had a greater amplitude in Global-KO animals as compared to PC-KO animals. Columns: mean, error bars: SEM. (**N**) Cumulative distributions for all recorded tremor bouts by strain. Inset in right panel highlights events with a duration between 5 and 10s.

To test if selective knockout of the GABA_Aα1_ subunit from Purkinje cells would be sufficient to induce a tremor phenotype, we performed similar tail-suspension recordings from PC-KO animals. As with Global-KO animals, a tremor phenotype was observed in PC-KO animals as compared to PC-WT and PC-HEMI animals, with a corresponding increase in power between 20Hz and 30Hz (PC-KO-Bl6 TM: 1.1 ± 0.08, n=18 animals; PC-WT-Bl6 TM: 0.65 ± 0.03, n=30 animals; PC-HEMI-Bl6 TM: 0.60 ± 0.04, n=10 animals; Black6 background TM across genotypes: p<0.0001, KW. PC-KO-Mixed TM: 1.2 ± 0.13, n=27 animals; PC-WT-Mixed TM: 0.76 ± 0.04, n=23 animals; PC-HEMI-Mixed TM: 0.76 ±0.04, n=22 animals; Mixed background TM across genotypes: p=0.0006, KW; Fig.3). Notably, a tremor phenotype was observed in PC-KO animals regardless of the background strain of the animal (PC-KO-Bl6 TM vs PC-KO-Mixed TM: p = 0.83, MW; Fig. 3J), indicating that background strain does not play a pivotal role in the expression of the tremor phenotype. There was also no significant difference in tremor severity between males and females (TM PC-KO-Bl6 males: 1.2 ± 0.11, n=7 animals, females: 0.9 ± 0.1, n=7 animals, p = 0.12, MW; TM PC-KO-Mixed males: 1.2 ± 0.17, n=15 animals, females: 1.2 ± 0.23, n = 12 animals, p = 0.45, MW, Fig. 3J). Together, these data indicate that selective knockout of the GABA_Aα1_ subunit from Purkinje cells is sufficient to induce a tremor, regardless of the background strain or sex of the animal.

Note that the frequency band of the tremor (20-30Hz) in PC-KO animals (Fig.3A-B) is similar to the tremor observed in global GABA_Aα1_ knockout animals (Fig. 3C) (51, 52), suggesting similar underlying mechanisms. However, the tremor was more prominent in the Global-KO mice (TM in KOs between lines: p<0.0001, KW; Global-KO vs PC-KO-Bl6 p < 0.0001, MW; Global-KO vs PC-KO-Mixed p < 0.0001, MW; Fig. 3J), suggesting that additional circuit elements or the timing of the knockout contribute to the overall phenotype in animals with a global knockout.

We observed that the tremor was not always present, but rather occurred in bouts. Visually, it appeared that PC-KO animals had fewer bouts of tremor than Global-KO animals, which may contribute to a lower overall TM value. To further explore the tremor phenotype, we quantified the rate of bouts, the length of bouts, and the amplitude of bouts (percent change in power) per animal, and compared across mouse lines (Fig. 3K-N). Global-KO mice displayed a higher rate of bouts (Global-KO: 18 ± 1 bouts/min, across mouse lines p < 0.0001, KW), longer bouts (Global-KO: 0.81 ± 0.08s; comparing across lines p < 0.0001, KW), *and* a greater amplitude of tremor per bout (Global-KO: 25 ± 1% change in force; comparing across lines p < 0.0001, KW) than PC-KO-Bl6 (rate, length, amplitude, p < 0.001 each, MWs) and PC-KO-Mixed animals (p < 0.001 each, MWs). This was apparent both in the mean values across animals (Fig.3K-M), and in the cumulative distributions of all recorded bouts (Fig.3N). There were no differences, however, between Purkinje cell specific knockout strains in the rate of tremor bouts (PC-KO-Bl6: 7.6 ± 0.8 bouts/min; PC-KO-Mixed: 6.2 ± 0.8 bouts/min, p = 0.12, MW), and only subtle differences in length (PC-KO-Bl6: 0.42 ± 0.03s; PC-KO-Mixed: 0.34 ± 0.02s, p=0.02, MW) and amplitude (PC-KO-Bl6: 18 ± 1 % change in force; PC-KO-Mixed: 16 ± 1% change in force, p = 0.02, MW; Fig. 3K-M), again suggesting that the background of the animal is not critical for expression of the tremor phenotype.

### Tremor in PC-KO animals is sensitive to Ethanol

Having determined that a selective knockout of the GABA_Aα1_ subunit from Purkinje cells is sufficient to induce a tremor phenotype, we next examined the pharmacology of this tremor -- specifically, if this tremor would be sensitive to ethanol (Figure 4). The tremor in Global-KO mice is sensitive to pharmacological agents including ethanol (7), similar to Essential Tremor in patients. We found that ethanol (1.25g/kg, i.p.) significantly reduced tremor in both PC-KO-Bl6 and PC-KO-Mixed animals (Bl6: p=0.005, Mixed: p=0.01, Friedmans; Fig. 4A), and the reduction in tremor largely persisted over the course of the hour examined, regardless of background (PC-KO-Bl6: TM reduced to 60 ± 5% 20minutes post injection, 68 ± 10% at 40minutes, and 76 ± 7% of baseline at 60minutes; PC-KO-Mixed: TM reduced to 61 ± 11% of baseline at 20 minutes post injection, 64 ± 12% at 40 minutes, 76 ± 15% at 60 minutes). In contrast, no reduction was seen following saline injection (PC-KO-Bl6: p = 0.36 Friedman, TM 130 ± 20%, 135 ± 23%, and 151 ± 24% of baseline at 20, 40, and 60 minutes post injection, respectively; PC-KO-Mixed: p=0.50 Friedman, TM 118 ± 29%, 105 ± 19%, and 147 ± 29% of baseline at 20, 40, and 60 minutes post injection, respectively).

**Figure 4.**
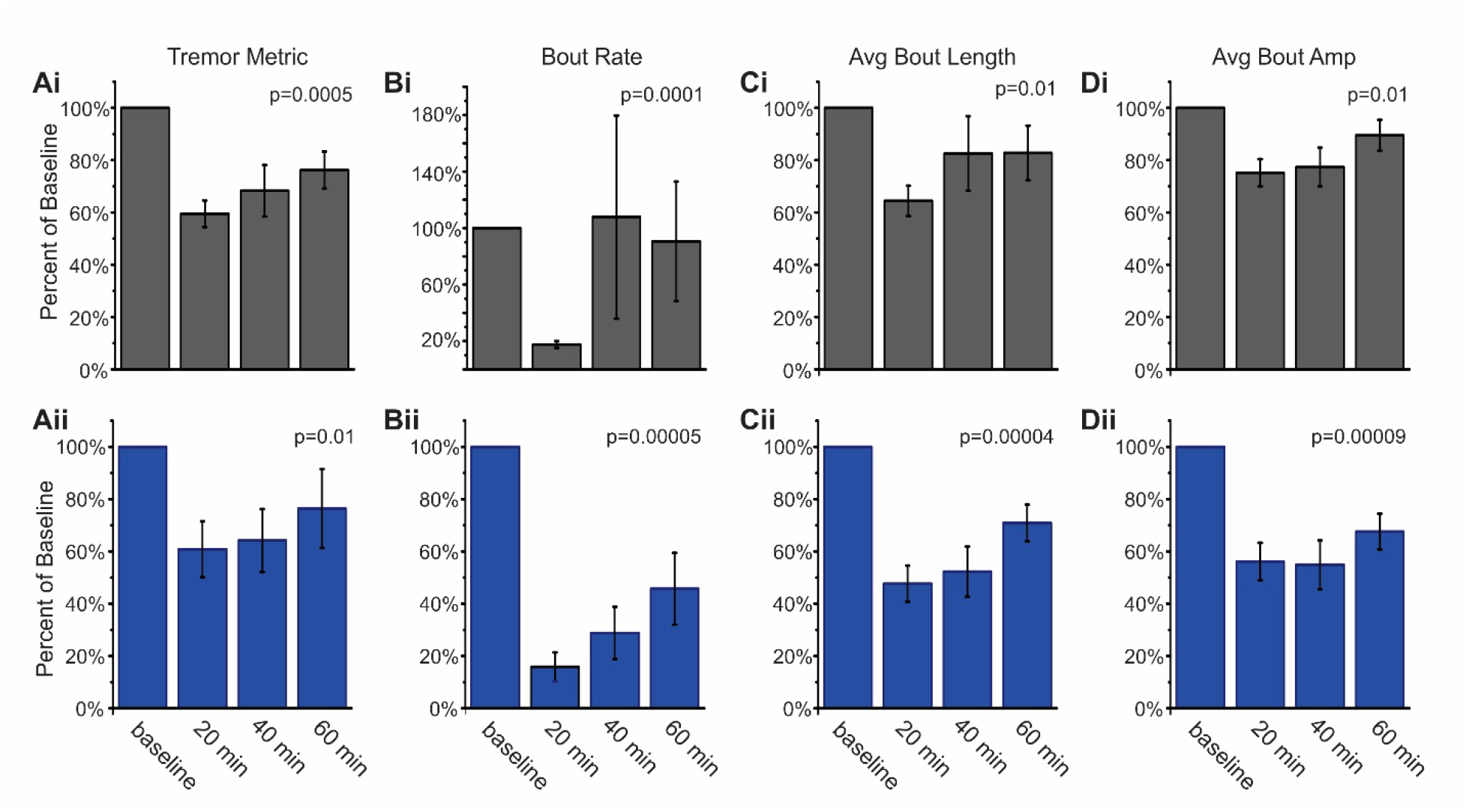
Ethanol inhibits tremor in PC-KO mice. Tremor Metric (**A**), rate of bouts (**B**), length of bouts (**C**), and amplitude of bouts (**D**) are all significantly reduced after i.p injection of 1.25 g/kg ethanol in PC-KO-Bl6 (**i, black**) and PC-KO-Mixed (**ii, blue**) animals. Percent of baseline (mean ± SEM) at 20, 40, and 60 minutes post-drug administration is shown for n=11 animals for each mouse line; p values from Friedman tests.

To determine if this decrease in the broad Tremor Metric was due to a reduced rate of tremor bouts, shorter tremor bouts, and/or a reduced amplitude per bout, we performed a more detailed analysis of the tremor bouts observed. Ethanol strongly reduced the rate of detected bouts (p < 0.001, Friedmans), especially at the earliest time point examined (Fig.4B). The length of bouts was also reduced, although this reduction was less dramatic than the initial reduction in the rate of bouts (Bl6: p = 0.01; Mixed: p < 0.01, Friedmans; Fig.4C). The amplitude of individual bouts was also reduced (Bl6: p = 0.01; Mixed: p < 0.01, Friedmans; Fig.4D), indicating a broad reduction in the rate, length, *and* strength of remaining bouts by ethanol. These data illustrate that tremor is strongly attenuated by ethanol administration, reminiscent of the global GABA_Aα1_ knockout model (7) and human Essential Tremor (8).

## DISCUSSION

To further investigate the tremor phenotype seen in mice with a global knockout of the GABA_Aα1_ subunit, we produced mice with a more restricted cerebellar Purkinje cell specific knockout of GABA_Aα1_. This resulted in an abolishment of all GABA_A_-mediated synaptic inhibition to Purkinje cells, but left GABA_A_-mediated inhibition to cerebellar molecular layer interneurons intact. As a result, any phenotype observed can be attributed, specifically, to a loss of inhibition to Purkinje cells. Purkinje cell specific knockout of GABA_Aα1_ did not result in reduced survival in animals, nor gross motor deficits as assessed by the accelerating rotarod test, but did result in a tremor phenotype. This tremor phenotype was less severe than tremor observed in Global-KO animals, but had similar properties (i.e., the tremor was focused in the 20-30Hz range). Hemizygous animals displayed GABA_A_ IPSCs in Purkinje cells with similar frequency, amplitude, and decay kinetics as wildtype animals, and, correspondingly, did not display a tremor phenotype. Finally, the tremor phenotype in PC-KO animals, similar to the tremor phenotype in global knockout animals (7), is sensitive to pharmacological inhibition through systemic administration of ethanol. In this regard, the tremor in PC-KO animals is reminiscent of the tremor seen in human Essential Tremor (8). These findings highlight a potentially key role in tremor for the cerebellum, Purkinje cells, and inhibition (and specifically, inhibition to Purkinje cells).

Previous studies using global GABA_Aα1_ knockout animals reported different phenotypes, with some studies reporting a tremor phenotype (7, 51, 52) but others finding instead increased mortality and an absence epilepsy phenotype (40). It was suggested that the background strain of the animals may underlie differences in the expressed phenotype (40). We therefore examined the impact of Purkinje cell specific knockout of GABA_Aα1_ on two different backgrounds. However, we found a similar tremor phenotype on both backgrounds, suggesting that the tremor phenotype is robust, and not heavily dependent on the background strain. We have not yet examined PC-KO animals for an epilepsy phenotype. Of interest, in addition to reports of epilepsy in mice with a loss of GABA_Aα1_, a recent study reported generalized epilepsy in zebrafish lacking GABA_Aα1_ (53). While it may seem unlikely that an epilepsy phenotype would be present in PC-KO animals, the engagement of the cerebellum during seizures has been repeatedly noted (43, 54–59). Additionally, mice with a loss of P/Q channels in the cerebellum display absence epilepsy (60, 61). In the future, it will therefore be interesting to examine if a cerebellar specific loss of the GABA_Aα1_ subunit can also produce an epilepsy phenotype.

While no decreased mortality was seen on either background in the PC-KO animals, we did find increased mortality in our Global-KO animals. We took care to have our Global-KO animals on a mixed background (matching previous studies which reported a tremor phenotype, but not increased mortality or epilepsy). Therefore, our findings suggest that background strain is not a likely explanation for differences in reported mortality. Early mortality does, however, appear to be a result of the global nature and/or timing of the knockout, as PC-KO animals do not display early mortality. The early mortality in Global-KO animals peaks around the time of weaning, is not reminiscent of human Essential Tremor, and provides experimental challenges. Early mortality is therefore not a desirable phenotype in a model of Essential Tremor, and PC-KO animals avoid this confound.

Perhaps the greatest drawback to the global knockout approach, however, is a lack of circuit insights. Given the hypothesized role of the cerebellum in tremor phenotypes (9, 15–17, 27), we were interested in testing the hypothesis that Purkinje cells play a key role in the observed tremor phenotype. Our findings, in which PC-KOs displayed a tremor phenotype reminiscent of the tremor in Global-KO animals, provides strong support for the role of the cerebellum, and Purkinje cells in particular, in tremor. Features of the tremor observed in the PC-KO animals mirror the global tremor sufficiently to suggest similar underlying mechanisms. However, the magnitude of the tremor was reduced in PC-KO animals, suggesting that intact inhibition to other circuit elements helps dampen the expression of tremor, such that it is amplified, and more frequent, in the Global-KO animals.

Specific knockout of the GABA_Aα1_ subunit from Purkinje cells resulted in a loss of GABA_A_-mediated inhibition to Purkinje cells, without compensatory expression of other α subunits. This indicates that a loss of inhibition, rather than altered kinetics of inhibition, to Purkinje cells is responsible for the tremor phenotype observed. Therefore, other manipulations to the circuitry that could similarly reduce inhibition to Purkinje cells (e.g. loss of molecular layer interneurons, or reduced drive to molecular layer interneurons) may also result in a tremor phenotype. The loss of GABAergic IPSCs in Purkinje cells in knockout animals further indicates that the tremor reducing effects of ethanol are unlikely to be mediated by an increase in GABAergic inhibition to Purkinje cells (e.g., through increased GABA release from molecular layer interneurons) (62–65). Either ethanol is working through a different mechanism to inhibit Purkinje cells, or is working upstream (e.g., to reduce excitatory drive to Purkinje cells) or downstream (e.g. in the cerebellar nuclei or further downstream).

While our findings provide some important insights, questions remain, including whether the loss of inhibition to Purkinje cells simply increases their net excitability or regularity of firing (66), or if the loss of inhibition (also) produces an increase in synchronization of Purkinje cells, and this specifically is what drives a tremor phenotype. Additionally, tremor is not constantly present in the animals, even when tail-suspended, but rather occurs in bouts. What serves as a trigger for tremor, switching the system into a pathological ‘tremor state’? Future work, using the Purkinje cell specific knockout animals developed here, can provide needed insight into these critical questions.

Future work can also examine the potential for non-tremor related phenotypes in PC-KO animals. Given the strength of the impact on inhibition to Purkinje cells (i.e., there was essentially an abolishment of IPSCs in Purkinje cells), it was notable that no effects were detected on the accelerating rotarod test. This suggests a minimal role of inhibition to Purkinje cells (or a compensatory mechanism) in learning and performance on the accelerating rotarod task. Results on the accelerating rotarod in previous studies in animals with a global knockout of GABA_Aα1_ have been mixed (50, 67). It will be of interest to see what phenotypes can, and cannot, be observed in a variety of cerebellar dependent tasks in PC-KO animals in future studies, as this could provide increased insight into the role of GABAergic inhibition to Purkinje cells (both from molecular layer interneurons and from other Purkinje cells) in cerebellar function (44, 66, 68–75). Similarly, the cerebellum is important for functions beyond motor control (76–81), and it will therefore also be of interest to examine potential non-motor phenotypes in PC-KO animals. That is, PC-KO animals provide a useful tool to study not only tremor, but also the role of the cerebellum, and in particular inhibition to Purkinje cells, in a range of conditions.

The pathophysiology of Essential Tremor largely remains a mystery. This underscores the need for models to study the disorder, but also makes finding great animal models especially challenging. The global GABA_Aα1_ knockout model provides a potential tool to study Essential Tremor, complementing the chemical (harmaline) model of tremor. The relevance of the global GABA_Aα1_ knockout tremor phenotype to Essential Tremor is not firmly established, but, importantly, the observed tremor phenotype is sensitive to both ethanol and propranolol (7). Minimally, therefore, the global GABA_Aα1_ knockout model provides a tool to examine this pharmacology. In this regard, it is important to note that the tremor observed in PC-KO animals is similarly sensitive to ethanol, while also benefiting from a more restricted loss GABA_Aα1_ (which further assists with interpretation of findings).

Our findings of a tremor phenotype in PC-KO animals highlights the relevance of Purkinje cells in tremor, illustrates that a deficit in inhibition can produce a tremor phenotype, and paves the way for future investigations into tremor, and the role of inhibition to Purkinje cells in cerebellar functions.

## Funding

This work was funded in part through a McKnight Land Grant Professorship award, the University of Minnesota’s MnDRIVE (Minnesota’s Discovery, Research and Innovation Economy) initiative, and an International Essential Tremor Foundation (IETF) grant (to EKM).

## Author contributions

AN collected and analyzed the electrophysiology data, with support from EKM. JK collected and analyzed the rotarod data, with support from AN and EKM. HG helped construct the tail suspension apparatus, AN and HG collected tremor data, and CS collected data examining the pharmacological responsiveness of tremor. CKM and EKM analyzed the tremor data. AN, JK, CKM, CS, and EKM wrote the paper. AN and EKM designed experiments and provided project management.

## Acknowledgements

We thank Michael Benneyworth and the University of Minnesota’s Mouse Behavior Core, for assistance with the accelerating rotarod testing, the other members of the Krook-Magnuson lab, especially Isaac Hoff for skilled colony management, and Stephen Fowler for assistance with establishing tremor measurements in our lab. This work was funded in part through a McKnight Land Grant Professorship award, the University of Minnesota’s MnDRIVE (Minnesota’s Discovery, Research and Innovation Economy) initiative, and an International Essential Tremor Foundation (IETF) grant (to EKM).

